# Individual differences in proprioception predict the extent of implicit sensorimotor adaptation

**DOI:** 10.1101/2020.10.03.324855

**Authors:** Jonathan S. Tsay, Hyosub E. Kim, Darius E. Parvin, Alissa R. Stover, Richard B. Ivry

**Author notes:** Corresponding author Information: Name: Jonathan Tsay, Address: 2121 Berkeley Way, Berkeley, CA 94704.

## Abstract

Recent studies have revealed an upper bound in motor adaptation, beyond which other learning systems may be recruited. The factors determining this upper bound are poorly understood. The multisensory integration hypothesis states that this limit arises from opposing responses to visual and proprioceptive feedback. As individuals adapt to a visual perturbation, they experience an increasing proprioceptive error in the opposite direction, and the upper bound is the point where these two error signals reach an equilibrium. Assuming that visual and proprioceptive feedback are weighted according to their variability, there should be a correlation between proprioceptive variability and the limits of adaptation. Alternatively, the proprioceptive realignment hypothesis states that the upper bound arises when the (biased) sensed hand position realigns with the target. When a visuo-proprioceptive discrepancy is introduced, the sensed hand position is biased towards the visual cursor and the adaptive system nullifies this discrepancy by driving the hand away from the target. This hypothesis predicts a correlation between the size of the proprioceptive shift and the upper bound of adaptation. We tested these two hypotheses by considering natural variation in proprioception and motor adaptation across individuals. We observed a modest, yet reliable correlation between the upper bound of adaptation with *both* proprioceptive measures (variability and shift). While these results do not favor one hypothesis over the other, they underscore the critical role of proprioception in sensorimotor adaptation, and moreover, motivate a novel perspective on how these proprioceptive constraints drive implicit changes in motor behavior.

**SIGNIFICANCE STATEMENT:** While the sensorimotor system uses sensory feedback to remain properly calibrated, this learning process is constrained, limited in the maximum degree of plasticity. The factors determining this limit remain elusive. Guided by two hypotheses concerning how visual and proprioceptive information are integrated, we show that individual differences in the upper bound of adaptation in response to a visual perturbation can be predicted by the bias and variability in proprioception. These results underscore the critical, but often neglected role of proprioception in human motor learning.

## INTRODUCTION

Accurate motor control requires the continuous calibration of the sensorimotor system, a process driven by the sensory feedback experienced over the course of movement. One of the primary learning processes involved in keeping the system calibrated is implicit sensorimotor adaptation (Shadmehr, Smith, & Krakauer, 2010; Taylor, Krakauer, & Ivry, 2014; Tseng, Diedrichsen, Krakauer, Shadmehr, & Bastian, 2007). Here, learning is assumed to be driven by sensory prediction error (SPE), the difference between the predicted feedback from a motor command and the actual sensory feedback.

Recent findings have shown that implicit adaptation in response to a visuomotor rotation (VMR) is remarkably invariant across a large range of error sizes and tasks (Kim, Morehead, Parvin, Moazzezi, & Ivry, 2018; Morehead, Taylor, Parvin, & Ivry, 2017). Even in response to large errors (e.g., 45°), the maximum amount of trial-to-trial change is around 1° - 2° (Bond & Taylor, 2015; Herzfeld, Vaswani, Marko, & Shadmehr, 2014; Kim et al., 2018; Morehead et al., 2017; Vandevoorde & Orban de Xivry, 2019; Wei & Körding, 2009) – not surprising for a system that likely evolved to adjust for subtle changes in the environment and body. More puzzling, the maximum degree of plasticity within this slow learning system is limited, reaching an asymptotic value of around 15° - 25° even after hundreds of trials (Bond & Taylor, 2015; Dang, Parvin, & Ivry, 2019; Haith, Huberdeau, & Krakauer, 2015; Kim et al., 2018; Morehead et al., 2017; Rand & Heuer, 2019; Werner et al., 2015) or across multiple test sessions (Stark-Inbar, Raza, Taylor, & Ivry, 2017; Wilterson & Taylor, 2019). As such, learning to compensate for large errors requires the recruitment of other learning processes such as explicit aiming strategies (Haith et al., 2015; Hegele & Heuer, 2010; Huberdeau, Haith, & Krakauer, 2015; McDougle & Taylor, 2019).

Although the mean upper bound for implicit adaptation to large visuomotor rotations averages around 20°, individual differences can be quite substantial. In standard VMR tasks, these differences are hard to detect during learning since participants eventually exhibit near-perfect performance, independent of the size of the perturbation. With these tasks, the individual differences become evident during the “washout” phase when feedback is eliminated, and participants are instructed to reach directly to the target (Bond & Taylor, 2015; Taylor et al., 2014). An alternative method is to use non-contingent, “clamped” visual feedback in which the angular trajectory of the feedback cursor is invariant, always following a path that is deviated from the target by a fixed angle (e.g., 15°). Despite instructions to ignore this feedback, the participants’ behavior reveals an automatic and implicit adaptation response, deviating across trials in the opposite direction of the clamp (Kim et al., 2018; Morehead et al., 2017; Tsay, Parvin, & Ivry, 2020). With this method, the error remains constant across trials; as such, the asymptote is not tied to changes in task performance (i.e., feedback terminating closer to the target), but rather, the asymptote reflects endogenous constraints. Across both methods (washout performance in tasks using contingent feedback or asymptotic performance in response to non-contingent feedback), the range of values is considerable. For example, in one study (Kim et al., 2018), the range of asymptotes in response to 15° clamped feedback was between 12° and 43° (mean = 18°, sd = 10°).

The factors which determine the upper bound of implicit adaptation are poorly understood. One hypothesis is that the limit reflects the interaction of visual and proprioceptive feedback. As adaptation progresses, the hand movements are adjusted away from the target, reducing the visual SPE (at least in standard VMR tasks). However, the change in hand direction away from the target results in an increase in a proprioceptive SPE, the difference between the expected and experienced signals of hand position. Importantly, the direction of the proprioceptive SPE is opposite to that of the visual SPE, and thus the response to these two SPEs are in the opposite directions. The asymptotic level of adaptation may thus reflect an equilibrium between learning from visual and proprioceptive error signals.

Studies of multisensory integration (Figure 1A) have shown that when participants estimate the location of their hand, they use a combination of visual and proprioceptive feedback, weighting each source based on their relative *reliability* (Burge, Ernst, & Banks, 2008; Ernst & Banks, 2002; Sober & Sabes, 2003; R. J. van Beers, Sittig, & Denier van der Gon, 1998; Robert J. van Beers, 2012; Robert J. van Beers, Wolpert, & Haggard, 2002). Consistent with this hypothesis, in the context of visuomotor adaptation, the response to a visual perturbation is reduced when noise is added to the visual feedback (Burge et al., 2008; Körding & Wolpert, 2004; Tsay, Avraham, et al., 2020; Robert J. van Beers, 2012; Wei & Körding, 2010). The corollary prediction, namely that the response to a visual perturbation should increase as a function of noise (i.e., variability) within the proprioceptive system, has not been tested.

**Figure 1:**
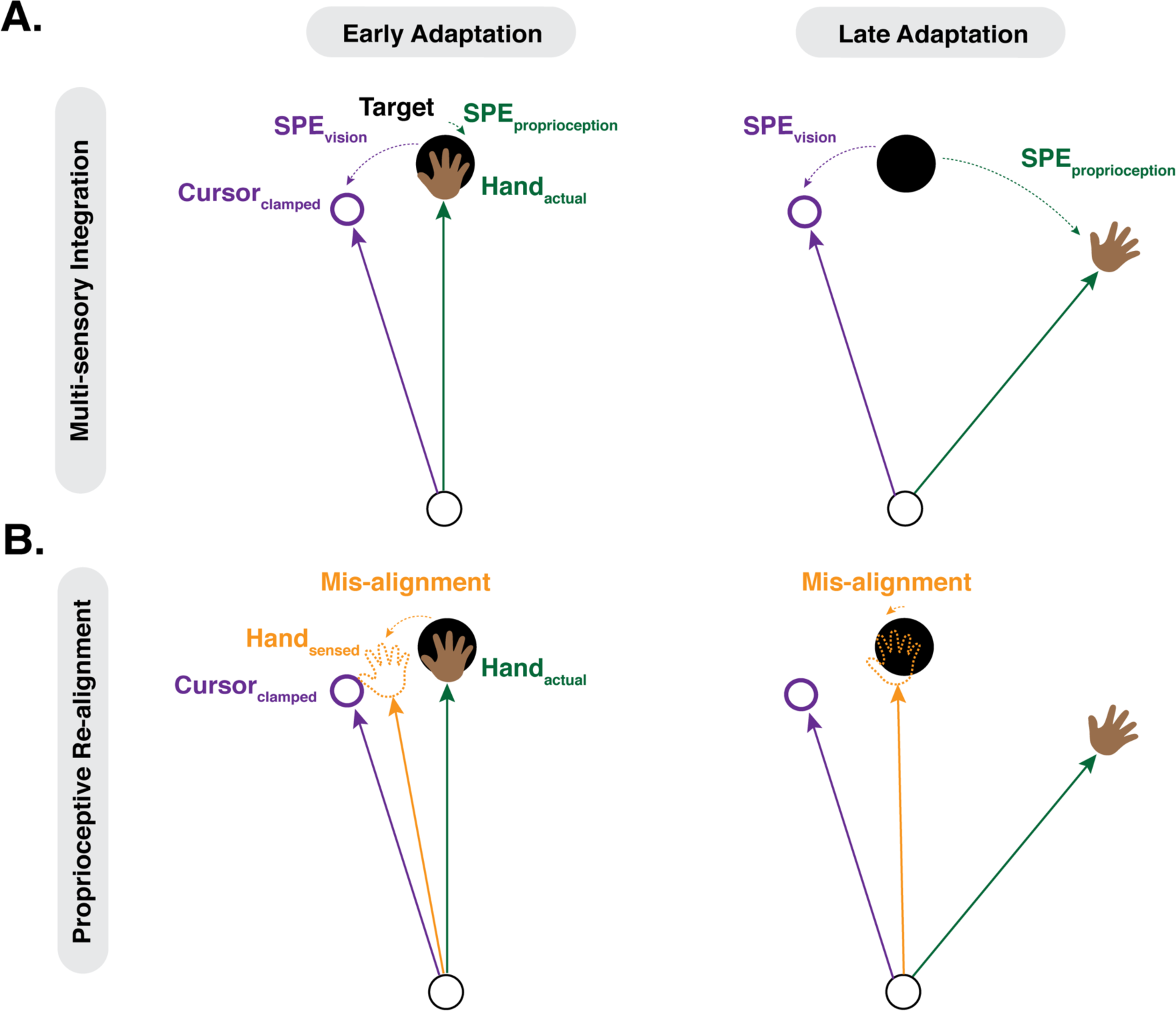
Two hypotheses concerning constraints on the upper bound of implicit adaptation. **A)** By the multisensory integration hypothesis, the upper bound of adaptation is the point of equilibrium between the visual SPE and the proprioceptive SPE. **B)** By the proprioceptive realignment hypothesis, the upper bound of adaptation occurs when the participant’s sensed hand position is at the target. Sensed hand position is a composite of visual-based inputs underlying the proprioceptive shift (target and cursor) and proprioception from the true hand position.

A second hypothesis relates to another way in which visual and proprioceptive information have been shown to interact during adaptation. The introduction of a visual perturbation creates a discrepancy between the visual and proprioceptive feedback. This discrepancy results in an immediate shift in the perceived location of the hand towards the visual feedback, a phenomenon referred to as a “proprioceptive shift.” The size of the shift tends to range between 5° - 10°, and remains relatively stable, evidenced by probing sensed hand position following passive hand displacement at various timepoints in an adaptation study (Cressman & Henriques, 2009, 2010; Ruttle, ‘t Hart, & Henriques, 2018). Similar to multisensory integration, this shift presumably reflects the operation of a system seeking to establish a unified percept from discrepant sensory signals (Cressman & Henriques, 2009, 2010; Rand & Heuer, 2019, 2020; Ruttle et al., 2018; Salomonczyk, Cressman, & Henriques, 2013; Synofzik, Thier, Leube, Schlotterbeck, & Lindner, 2010; Robert J. van Beers et al., 2002; Wilke, Synofzik, & Lindner, 2013; Zbib, Henriques, & Cressman, 2016).

The processes underlying proprioceptive shift may also contribute to the upper bound of implicit adaptation (Figure 1B). This shift introduces a different error signal, the discrepancy between the target and the sensed hand position (i.e. the difference between the expected trajectory to the target and the trajectory towards the perceived hand position). A learning process seeking to nullify this error signal would also drive the hand direction away from the perturbation (i.e., the opposite direction of the proprioceptive shift). By this view, implicit adaptation would reach an asymptote when the sensed hand position is “re-aligned” with the target, and as such, the asymptote would correlate with the size of the proprioceptive shift: A larger deviation in hand angle would be required to nullify a large proprioceptive shift. This prediction is consistent with the results of a recent VMR study (Ruttle, Hart, & Henriques, 2020).

To examine these two hypotheses in tandem, we exploit here natural variation across individuals, examining the relationship between individual differences in proprioceptive variability and proprioceptive shift with the upper bound of implicit adaptation. To operationalize our proprioceptive measures, participants were asked to report the position of their hand after passive displacement. These proprioceptive probes were obtained before, during, and after an extended block of trials in which the visual feedback was perturbed. From these data, we could use standard psychophysical methods to estimate for each participant, the bias and variability in their sense of proprioception, with the bias providing an assay of proprioceptive shift. In Experiment 1, the upper bound on implicit adaptation was estimated by measuring the participants’ aftereffect in response to a response-contingent visuomotor rotation. In Experiment 2, the upper bound was estimated using the asymptotic response to clamped visual feedback.

## METHODS

### Participants

Undergraduate students were recruited from the UC Berkeley community (N = 62; age = 18 – 22; 45 women, 17 men) and either received course credit or financial compensation for their participation. As assessed by the Edinburgh handedness inventory, all of the participants were right handed (Oldfield, 1971). The protocol was approved by the IRB at UC Berkeley.

### Experimental overview

Each experiment involved a mix of reaching trials and proprioceptive probe trials. For both tasks, the participants were seated in front of a custom table top setup and placed their hand on a digitizing graphics tablet (49.3 cm by 32.7 cm, Intuos 4XL; Wacom, Vancouver, WA, sampling rate = 200 Hz.) that was horizontally aligned with and positioned below an LCD monitor (53.2 cm by 30 cm, ASUS). The participant’s view of their hand was occluded by the monitor, and the room lights were extinguished to minimize peripheral vision of the arm. On reaching trials, arm movements were made by sliding a digitizing pen, embedded in a custom handle, across the table. On proprioceptive trials, the participant held the digitizing pen and the experimenter moved the participant’s arm.

### Reaching Trials

Reaches were made from a start location to a target, located at various locations (see below). The start location was indicated by a white ring (6 mm diameter) and the target by a blue circle (6 mm diameter), with the radial distance between the start location and target fixed at 16 cm. To initiate a trial, the participant moved her hand to the start location. Visual feedback of the hand position was given via a cursor (white circle 3.5 mm diameter) only when the hand was within 1 cm of the start position. Once the hand remained within the start location for 500 ms, the target appeared, serving as a cue to indicate the location of the target and an imperative to initiate the reach. To discourage on-line corrections, participants were instructed to perform ‘shooting’ movements, making a rapid movement that intersected the target.

There were two types of feedback trials: Veridical and perturbed. On veridical trials, the cursor corresponded to the position of the hand. On perturbation trials, the cursor was either rotated relative to the hand position (visuomotor rotation, Exp 1) or restricted to an invariant path along a constant angle with respect to the target (visual clamp, Exp 2). On feedback trials, the radial position of the cursor matched the radial position of the hand until the movement amplitude reached 16 cm (the radial distance of the target), at which point the cursor froze. On no-feedback trials, the cursor was blanked when the target appeared, and did not re-appear until the participant had completed the reach and returned to the start location for the next trial.

Movement time was defined as the interval between when the hand movement exceeded 1 cm from the start position to when the radial distance of the movement reached 16 cm. To ensure that the movements were made quickly, the computer played a prerecorded message “too slow” if movement time exceeded 300 ms. If the movement time was less than 300 ms, a neutral ‘knock’ sound was generated, informing the participant that the reach speed had fallen in the acceptable window. There were no constraints on reaction time.

### Proprioceptive Probe Trials

To probe proprioceptive variability, the experimenter sat at the opposite side of the table, across from the participant. From this position, the experimenter could passively move the participant’s right hand to different probe locations (see below). The participant was instructed to hold the digitizing pen, but to maintain a passive state, one that allowed the experimenter to move the participant’s right hand with minimal resistance. To produce the passive movements, the experimenter used her left hand to move the participant’s right hand, maintaining contact throughout the proprioceptive probe block.

The experimenter initiated each trial by moving the participant’s hand into the start position, at which point the word ‘Ready’ appeared on the screen. The experimenter then hit the space bar with her right hand, at which point the word ‘Ready’ disappeared and a number specifying the desired target location appeared on the corner of the monitor closest to the experimenter (Figure 2). A small cloth cover was placed at this corner to prevent the participant from seeing the number. The experimenter moved the participant’s hand to the specified target location. Once the participant’s hand was at the target location (2 cm diameter tolerance window), the word ‘Ready’ again appeared and the experimenter hit the space bar to advance the trial. A filled white circle (3.5 mm diameter) then appeared at a random position on the monitor. The participant used her left hand to move a mouse (Logitech Trackman Marble), positioning the cursor above the sensed position of their right hand. When satisfied with the position of the cursor, the participant clicked the mouse button. The participant was allowed to modify their response by repositioning the mouse and clicking again. When the participant confirmed that the trial was complete, the experimenter hit the space bar, at which point the cursor disappeared. The experimenter then moved the participant’s hand back to the start position to initiate the next trial. The start position remained on the screen for the duration of the proprioceptive probe trials.

**Figure 2:**
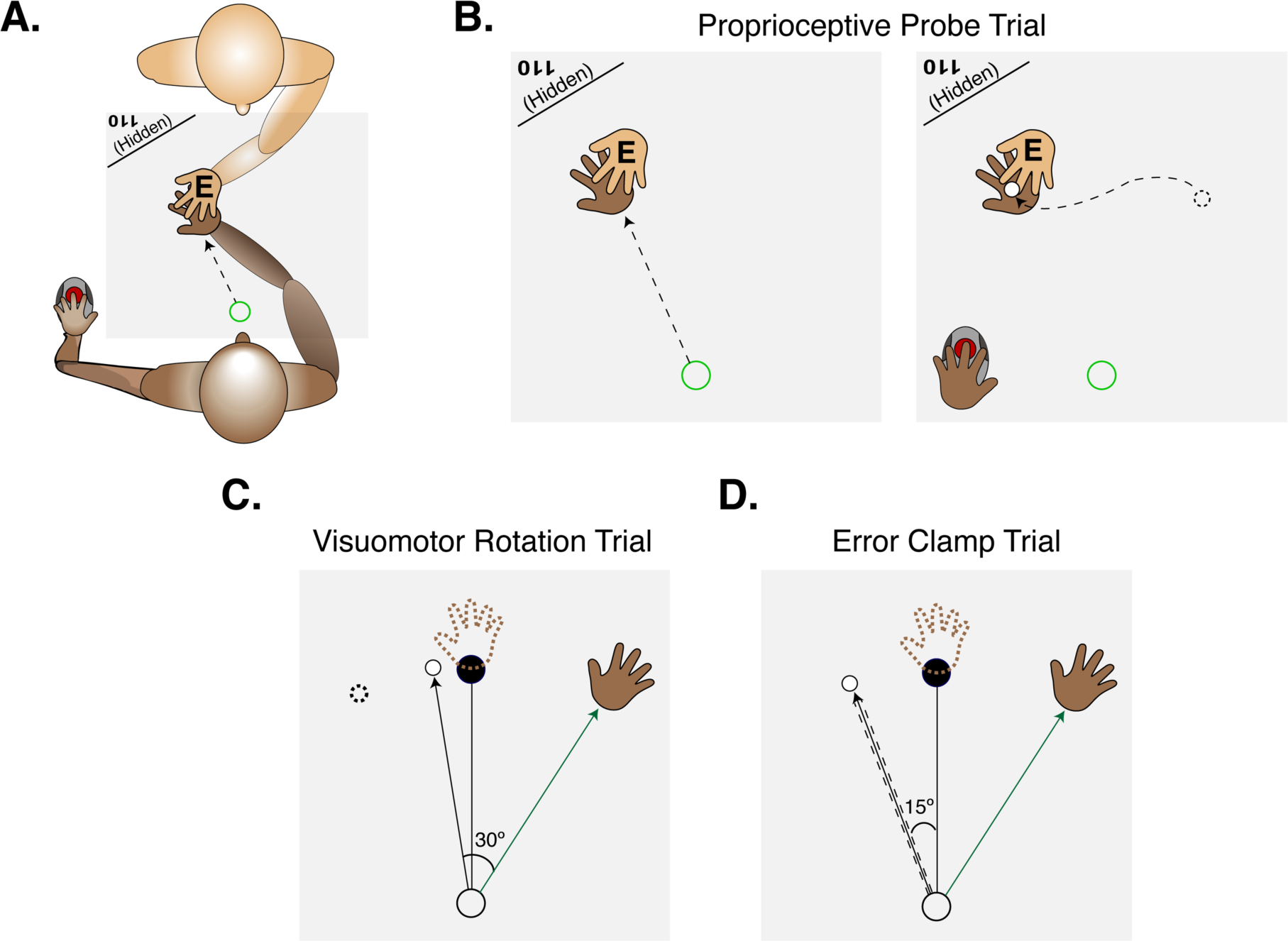
Experimental overview. **A)** Experimental setup for proprioceptive probe trials. The experimenter (top, with their hand labeled with an ‘E’) sat opposite the participant (bottom) and moved their hand from the start position to a specified location. The location (e.g. 110°) was signaled to the experimenter via text which appeared on the corner of the horizontal monitor, behind a cloth which prevented the participant from seeing the text. **B)** After the participant’s hand was passively moved to the probe location, a cursor appeared at a random position on the screen. The participant used their left hand to move the cursor to the sensed hand position. **C)** In Exp 1, a rotation was applied to the cursor. The task error introduced by the rotation is nullified if the participant moves in the opposite direction of the rotation. **D)** In Experiment 2, the cursor was clamped, independent of hand position. Participants were told to ignore the error clamp and aim straight for the target. The depicted trials in Panels C and D provide examples of performance late in the adaptation block.

We opted to use a tolerance window of 2 cm in positioning the hand, a value that was large enough for the experimenter to guide the participant’s hand to the target location without feedback, but also small enough to ensure minimal variation in target positions across trials. Note that variance in the position of the hand was irrelevant given that the proprioceptive judgments were recorded as the difference from the perceived location of the hand (mouse click) and the actual position of the hand.

### Experiment 1, Movement-contingent, rotated feedback

Reaching and proprioceptive trials were performed to 5 targets located within a wedge (at 70°, 80°, 90°, 100°, 110°, with 90° corresponding to straight ahead). The trials were arranged in cycles of one trial per target, with the order randomized within a cycle.

The experiment began with a brief phase to familiarize the participants with the reaching task. This consisted of 10 baseline reaching trials in which no visual feedback was provided, followed by 10 baseline trials with online, veridical feedback. The latter was used to emphasize that the movement should be produced to shoot through the target and demonstrate that the feedback would disappear once the movement amplitude exceeded the radial distance of the target.

The participant then completed a block of 50 baseline proprioceptive probe trials. Following this, the reaching task resumed but now the feedback perturbed. To minimize awareness of the perturbed feedback, the angular deviation of the cursor was increased in small, incremental steps of 0.33° per trial, reaching a maximum of 30° after 90 trials. Across participants, we counterbalanced the direction of the rotation (clockwise or counterclockwise).

Following the initial 90 perturbation trials, the participant then completed 7 more blocks, alternating between proprioceptive probe trials (30 per block) and reaching trials (40 per block, at the full 30° rotation). With this alternating schedule, we sought to obtain stable measures of proprioception following adaptation, while minimizing the effect of temporal decay on adaptation. These blocks were intermixed with four blocks of 5 no-feedback trials with instructions to reach directly to the target despite the absence of feedback. These no-feedback blocks occurred after the first gradual perturbation block, the second fixed perturbation block, the third perturbation block, and the fourth proprioceptive probe block. These no-feedback trials provided the primary data for our measure of adaptation. By having four of these probes, we were also able to assess the time course of adaptation. To complete the session, the participants completed 50 reaching trials with veridical feedback to ensure that the residual effects of adaptation were removed.

Each participant returned for a second session, 2 to 14 days after the first session. The experimental protocol was identical on day 2, allowing us to assess test-retest reliability of the various measures of adaptation and proprioception.

### Experiment 2, non-contingent, clamped feedback

The key change in Exp2 was the use of the visual clamp method during the perturbation trials. This form of feedback has been shown to produce robust adaptation with minimal awareness (Kim et al., 2018; Morehead et al., 2017; Jonathan S. Tsay, Parvin, & Ivry, 2020). Moreover, adaptation with this method will reach an upper bound that is not constrained by performance error (e.g., distance between cursor and target which is reduced over time with contingent feedback as in Exp 1), but presumably reflects factors intrinsic to each participant. Based on previous work, we expected to observe a broad range of upper bounds across our sample, a desirable feature to examine individual differences.

The basic method for the reaching and proprioceptive probe trials was similar to that used in Exp 1 with a few changes. First, we used a finer sampling of the workspace for the proprioception task, with target locations spaced every 5° (70°, 75°, 80°, 85°, 90°, 95°, 100°, 105°, 110°). Although participants were not explicitly queried in Exp 1, we were concerned that some participants may have noticed that there were only five discrete target locations, which could potentially bias their responses; that is, the proprioceptive reports might be based on their memory of a previously reported hand position rather than relying solely on the current proprioceptive signal. The finer sampling should reduce the utility of memory-based reports. Second, for the reaching task, we opted to keep the spacing as in Exp 1 (10° apart) but increased the size of the wedge, with the target locations spanning the range of 50° - 130°. This change was motivated by pilot work suggesting that adaptation to a visual clamp is more consistent when the movements are made in a larger workspace. Note that it was necessary to limit reaching in one direction, away from the body, given the workspace limitations imposed by the tablet and our decision to have the movement amplitude be 16 cm.

We also modified the block structure. Experiment 2 began with a proprioception block (one cycle, 1 trial per 9 targets) to familiarize the participant with this task. The participants then completed a block of reaching trials without visual feedback (9 targets, 27 trials total), followed by a block of reaching trials with veridical feedback (72 trials) and another proprioception block (72 trials, with a break after 36 trials). The participant then completed the perturbation block, composed of 180 trials (break after the first 90). For these trials, the cursor always followed a 16 cm straight trajectory offset by 15° from the target (clockwise or counterclockwise, counterbalanced across participants). The radial distance of the cursor, relative to the start position, was yoked to the participant’s hand. Thus, the motion of the cursor was temporally correlated with the participant’s hand, but its direction was fixed, independent of the angular position of the participant’s hand. Just before the start of this block, the error clamp was described to the participant and she was told to ignore this “feedback” signal, always attempting to reach directly to the target. To help the participant understand the invariant nature of the clamp, three demonstration trials were provided. On all three, the target appeared straight ahead at 90° and the participant was told to reach to the left (demo 1), to the right (demo 2), and backward (demo 3). On all three of these demonstration trials, the cursor moved in a straight line, 15° offset from the target. In this way, the participant could see that the spatial trajectory of the cursor was unrelated to their own reach direction.

Following the initial 90 trials with clamped feedback, the participant completed seven blocks, alternating between the proprioception task (36 trials/block, four blocks) and the reaching task with clamped feedback (90 trials/block, three blocks).

Given the impressive reliability results from Exp 1 (see below), we limited testing to a single session.

### Data Analysis

The experimental software and analyses were performed using custom scripts in Matlab and R.

The evaluation of our core hypotheses involves three variables of interest: Implicit adaptation, proprioceptive shift, and proprioceptive variability. The dependent variable for implicit adaptation was the change in hand angle from baseline, where hand angle was defined as the signed angular difference between the position of the hand at peak velocity and target, relative to the start location. In Exp 1, the measure of implicit adaptation was the hand angle during the no-feedback aftereffect trials (blocks 2 and 3, since adaptation saturated at block 2). In Exp 2, we used the mean hand angle during the last three blocks (block 2 - 4) of the error clamp trials since adaptation had reached a stable asymptote by block 2. For both experiments, the adaptation analyses were performed after correcting for any bias observed during the last two baseline cycles (Exp 1: 10 trials; Exp 2: 18 trials). Trials in which the hand angle exceeded three standard deviations from a moving 5-trial average were excluded from the analyses (Exp 1: 1.2% ± 0.6% per participant; Exp 2: 0.5% ± 0.3% per participant).

For proprioception, we recorded the x and y coordinate of each hand location report and calculated the Euclidean distance of this location to the actual hand location. From these data, we calculated the mean sensed hand position, relative to the target for each block of proprioceptive reports. Proprioceptive shift is the difference between the mean sensed hand position for each proprioceptive report block and the mean sensed hand position on the baseline block. For each block, we also calculated the standard deviation of the proprioceptive reports for each block, our measure of proprioceptive variability.

Exp 1 dependent measures were entered into a linear mixed effect model (R function: lmer), with Block and Day as fixed factors, and Participant ID as a random factor. Exp 2 dependent measures were entered into a linear mixed effect model, with Block as the only fixed factor and Participant ID as the random factor. All post-hoc t-tests were two-tailed, and Bonferroni corrected for multiple comparisons. Standard effect sizes are reported (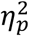 for fixed factors; Cohen’s *d*_*z*_ for within-subjects t-tests) (Lakens, 2013).

## RESULTS

### Experiment 1

The main goal of Exp 1 was to examine the relationship between implicit adaptation and individual differences in proprioception (proprioceptive shift and proprioceptive variability). For implicit adaptation, we focus on the change in heading angle on trials without feedback (aftereffect) following exposure to a 30° rotation of the visual feedback. Since the perturbation was introduced in a gradual manner, we assume the resulting recalibration of the sensorimotor map was implicit.

#### Implicit Adaptation

In order to track the time course of implicit adaptation, we measured mean hand angle during four no-feedback blocks, one at the end of the baseline block and three during the adaptation phase. There was a large main effect of block (Figure 3B) (*F*_4,261_ = 93.0 *p* < 0.001, *η*^2^ = 0.85), with the mean hand angles in each no-feedback block significantly different from baseline (*all t*_261_ > 24.1, *p*_*bf*_ < 0.001, *d*_*z*_ > 4.4). The mean hand angle increased from aftereffect block 1 to aftereffect block 2 (Figure 2A, *t*_261_ = 5.5, *p*_*bf*_ < 0.001, *d*_*z*_ = 1.0). There was no difference between the means in the second and third aftereffect blocks (*t*_261_ = 0.5, *p*_*bf*_ = 1, *d*_*z*_ = 0.01), suggesting that implicit adaptation in response to a 30° rotation saturated between 22° - 26°. The mean hand angle in the fourth aftereffect block was significantly lower than the third aftereffect block (*t*_261_ = –6.4, *p*_*bf*_ < 0.001, *d*_*z*_ = –1.2). Given that this block occurs after a set of proprioceptive probe trials, the difference here may indicate that proprioceptive trials had an attenuating effect on implicit adaptation (‘t Hart & Henriques, 2016).

**Figure 3:**
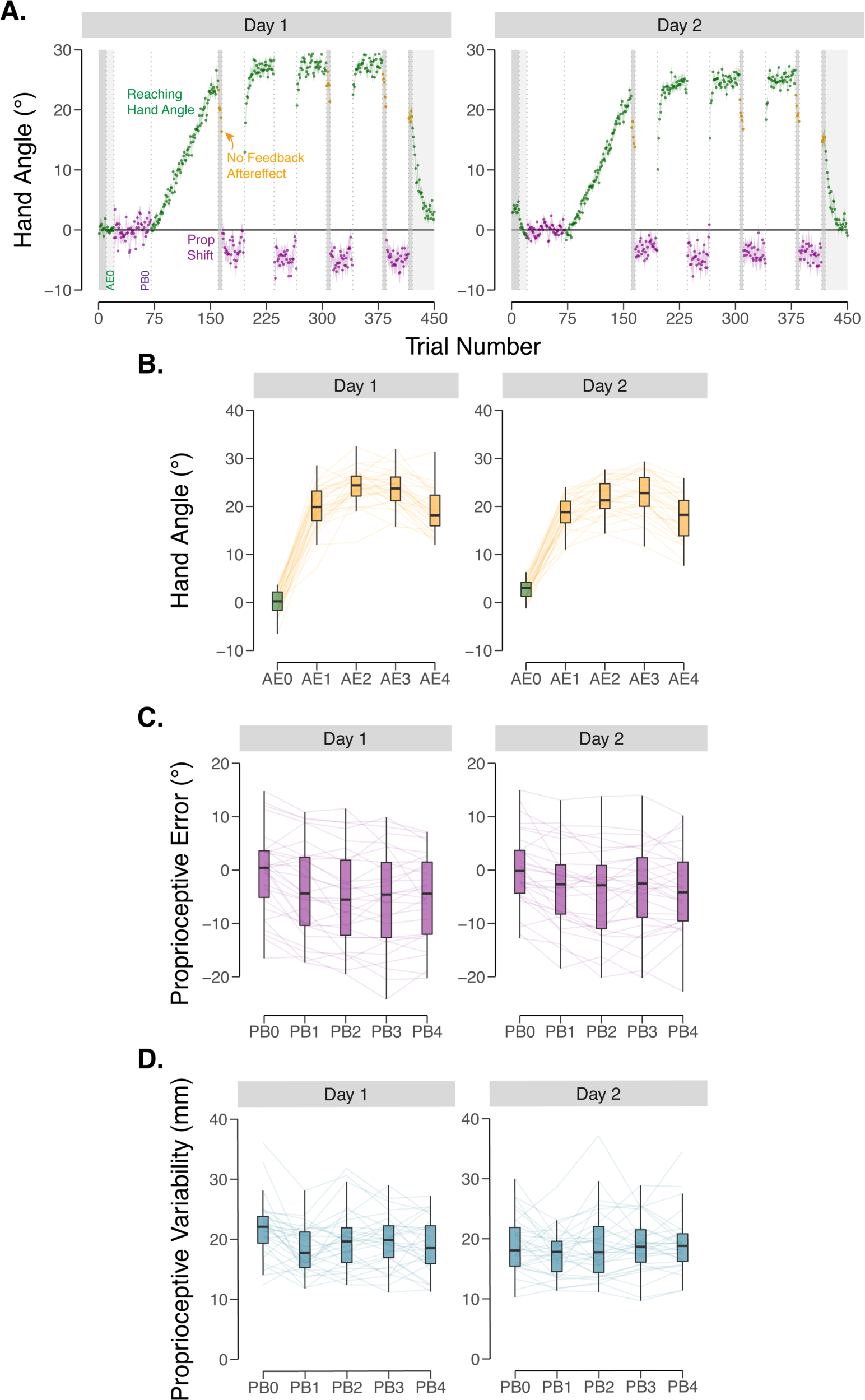
Performance on adaptation and proprioception probe tasks in experiment 1. **A)** Group means across test session (Left – Day 1, Right – Day 2). After a period of baseline trials, participants were exposed to a gradually increasing visuomotor rotation up to 30°, where it was then held constant. Participants performed blocks of visuomotor rotation trials (hand angle shown in green), no feedback aftereffect trials (hand angle shown in yellow), and proprioceptive probe trials (shift in perceived position shown in purple). Vertical dotted lines indicate block breaks. Shaded trials indicate reaching trials either with no feedback (dark grey) or with veridical feedback (light grey). Shaded regions indicate ± SEM. Baseline blocks for reaching hand angle (AE0) and proprioceptive probes (PB0) are labeled. **B)** Hand angle during no feedback aftereffect blocks. **C)** Proprioceptive errors for each proprioceptive block. **D)** Variability of proprioceptive judgments for each proprioceptive probe block. Thin lines indicate individual subjects. Box plots indicate min, max, and the 1^st^/3^rd^ interquartile range.

We next assessed whether adaptation remained stable across days: there was no main effect of Day, indicating that adaptation remained similar in both sessions (*F*_1,261_ = 0, *p* = 1, *η*^2^ = 0.04); however Day interacted with Block (*F*_4,261_ = 6.3, *p* < 0.001, *η*^2^ = 0.01). Post-hoc t-tests revealed that the aftereffect was smaller on day 2 compared to day 1 in block 3 (*t*_261_ = –5.1, *p*_*bf*_ < 0.001, *d*_*z*_ = 0.91). A similar pattern was evident in the other blocks, with the magnitude of the aftereffect lower on day 2 by about 4°. This attenuation has been observed in previous studies (Avraham, Ryan Morehead, Kim, & Ivry, 2020; Leow, Marinovic, de Rugy, & Carroll, 2020; Stark-Inbar et al., 2017; Wilterson & Taylor, 2019).

#### Proprioceptive Shift

We then assessed whether the exposure to the rotation resulted in a proprioceptive shift, quantified as the angular change in the centroid of proprioceptive estimates, relative to the baseline. The effect of Block was significant (Figure 3C) (*F*_4,261_ = 4, *p* = 0.003, *η*^2^ = 0.27), with a ∼4° proprioceptive shift towards the rotated feedback from baseline to PB1 (*t*_261_ = – 3.9, *p*_*bf*_ = 0.005, *d*_*z*_ < –0.7). Consistent with a previous study (Ruttle, Cressman, ‘t Hart, & Henriques, 2016), the shift remained stable across successive blocks (all pairwise comparisons of successive blocks in Day 1 were not significant: *t*_261_ < 0.1, *p*_*bf*_ = 1, *d*_*z*_ < 0.03). In addition, the magnitude of the proprioceptive shift was stable across days, with neither the effect of Day (*F*_1,261_ = 0, *p* = 1, *η*^2^ = 0.004), or significant Day x Block interaction (*F*_4,261_ = 0.3, *p* = 0.84, *η*^2^ = 0.004), consistent with the findings of a previous study (Liu, Sexton, & Block, 2018).

#### Proprioceptive Variability

To operationalize proprioceptive variability, we determined the Euclidian distance (x,y) from each hand report response for a given block to the average x and y coordinates of the reports for that block (Figure 3D). There was a main effect of block (*F*_4,261_ = 2.8, *p* = 0.02, *η*^2^ = 0.06), a modest effect that was driven by a decrease in proprioceptive variability from baseline (PB0) to PB1 (*t*_261_ = –4.0, *p*_*bf*_ < 0.001, *d*_*z*_ = –0.7). Given that this effect is most pronounced on Day 1, it may reflect the participants’ increased familiarity with the task (Liu et al., 2018). Nonetheless, proprioceptive variability was relatively stable across blocks (all remaining pairwise comparisons: *t*_261_ < 2.4, *p*_*bf*_ > 0.16, *d*_*z*_ < 0.4).

Proprioceptive variability attenuated between day 1 and day 2. There was a main effect of Day (*F*_1,261_ = 9.6, *p* = 0.002, *η*^2^ = 0.02), with proprioceptive variability being overall lower in day 2, consistent with the notion that increased familiarity with the task led to more consistent proprioceptive judgments. Post-hoc t-tests show that this effect was primarily driven by differences between proprioceptive variability in day 1 compared to the baseline proprioceptive variability in day 2 (Day 2 baseline vs Day 1 PB1 – PB4; *all t*_261_ < 3.9, *p*_*bf*_ < 0.001, *d*_*z*_ < –0.7). Previous studies comparing proprioceptive variability between days have also observed a similar improvement (Avraham et al., 2020). The overall pattern was nonetheless similar across both days, with no interaction observed between Block and Day (*F*_4,261_ = 1.5, *p* = 0.21, *η*^2^ = 0.02).

#### Reliability of the dependent variables

Analyses which involve correlating individual differences across different measures are limited by the reliability of each measure. Thus, before turning to the correlational analyses between the proprioceptive measures and implicit adaptation, we assessed the reliability of our core measures across sessions. For adaptation, we used the mean of the last two aftereffect blocks (AE2 - AE3) given that adaptation has reached its limit by these blocks. For proprioceptive shift, we used the mean proprioceptive shift of all blocks (PB1 – PB4) after the perturbation was introduced relative to baseline. For proprioceptive reliability, we used the proprioceptive variability from all blocks (PB0 – PB4). The between-session correlations were significant for all three dependent variables (Figure 4A-C) (implicit adaptation: *R* = 0.53, *p* = 0.002; proprioceptive variability: *R* = 0.72, *p* < 0.001; proprioceptive shift: *R* = 0.52, *p* < 0.001), indicating that the individual differences were reasonably stable.

**Figure 4:**
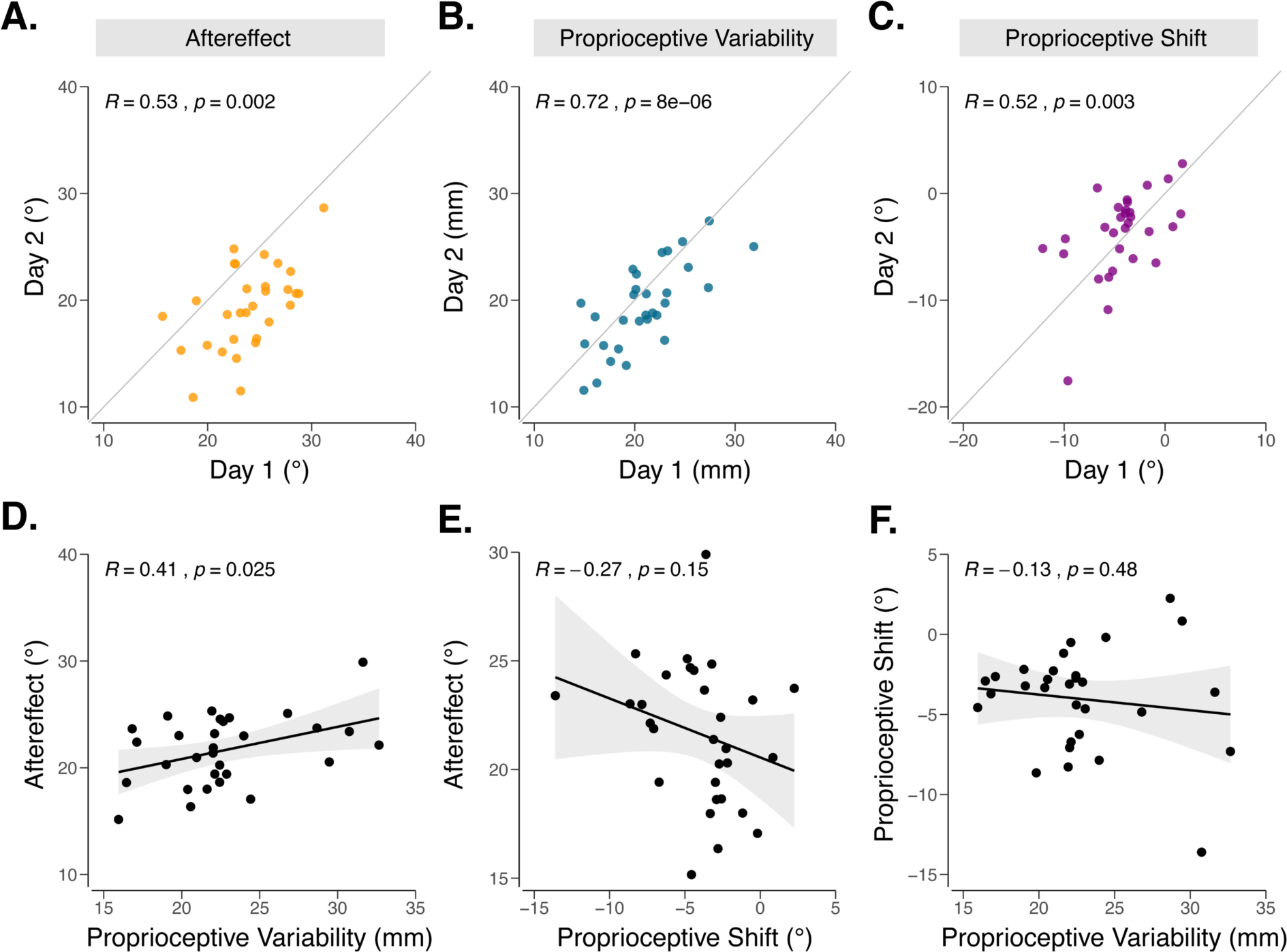
Inter-individual differences analyses in experiment 1. Test-retest reliability, measured across days for **A)** aftereffect from adaptation (yellow) **B)** proprioceptive variability (blue) and **C)** proprioceptive shift (purple). Correlations between different dependent variables: **D)** proprioceptive variability vs aftereffect; **E)** proprioceptive shift vs aftereffect; **F)** proprioceptive variability vs proprioceptive shift. Black line denotes the best fit regresion line and the shaded region indicates the 95% confidence interval.

#### Correlating Adaptation and Proprioception

Having established that these dependent variables were reliable across days, we next asked whether differences in implicit adaptation could be accounted for by individual differences in proprioception. We opted to focus on aftereffects on day 1 data given the evidence that adaptation can change across sessions, either from interference (Avraham et al., 2020) or savings (Yin & Wei, 2020), with our own data showing the former pattern. For a more stable measure of proprioceptive shift and proprioceptive variability, we averaged the data from both days.

According to the multisensory integration hypothesis, we should expect a positive correlation between proprioceptive variability and the extent of adaptation since, all other things being equal, noisier proprioception would diminish the relative weighting given the proprioceptive sensory prediction error. Consistent with this prediction, the two measures were positively correlated (Figure 4D: *R* = 0.41, *p* = 0.025).

According to the proprioceptive realignment hypothesis, we should expect a correlation between the proprioceptive shift and implicit adaptation. Given that these two effects should be in opposite directions, the correlation should be negative: A larger (more negative) proprioceptive shift from would require a larger change in hand angle for the hand to be perceived at the target location. Although the pattern was in the predicted direction, the correlation was not significant (Figure 4E: *R* = 0.27, *p* = 0.15).

We also examined the correlation between proprioceptive shift and proprioceptive variability. Although we had no strong *a priori* expectations here, a signal-dependent perspective might predict a negative correlation (Harris & Wolpert, 1998): Larger shifts would be more variable. Similarly, one might suppose that the perceived location of the hand might be more malleable if the inputs are more variable. However, proprioceptive variability and proprioceptive shift were not correlated (Figure 4F: *R* = –0.13, *p* = 0.48), an observation in line with a previous study looking at the (absent) relationship between the magnitude of cross-sensory calibration and signal reliability (Burge et al., 2008; Wei & Körding, 2010).

### Experiment 2

Exp 2 (n = 32) provided a second test of the multisensory integration and proprioceptive realignment hypotheses, using a visual error clamp in which the feedback cursor was always offset from the target by 15°. Compared to Exp 1 where the contingent feedback constrained the degree of adaptation, we expected the clamp to yield a greater range of values for implicit adaptation.

#### Implicit Adaptation

The participants’ reaches shifted in the opposite direction of the error clamp feedback, the signature of implicit adaptation (Figure 5). The hand angle data (CB0 – CB4, using the last 90 trials of CB1, and all 90 trials in CB2 – CB4) showed a main effect of Block (*F*_4,124_ = 55.1, *p* < 0.01, *η*^2^ = 0.06), with post-hoc comparisons indicating that the mean hand angle in each block was significantly greater than baseline (all *t*_124_ > 10.75, *p* < 0.001, *d*_*z*_ > 1.9). The mean values were not significantly different from one another for the four clamp blocks (all pairwise t-tests: *t*_124_ < 2.47, *p* > 0.13, *d*_*z*_ < 0.44), indicating that participants had reached the asymptote of adaptation by the end of the first clamp block. To obtain a single measure of adaptation for each participant, we took the mean hand angle over the last three clamped feedback blocks. The mean change in hand angle was 17.5° ± 13.9°. As expected, the range of asymptotic values was considerably larger in Exp 2 (range = -6.5° – 58.5°) compared to Exp 1 (range = 13.5° – 33.9°).

**Figure 5.**
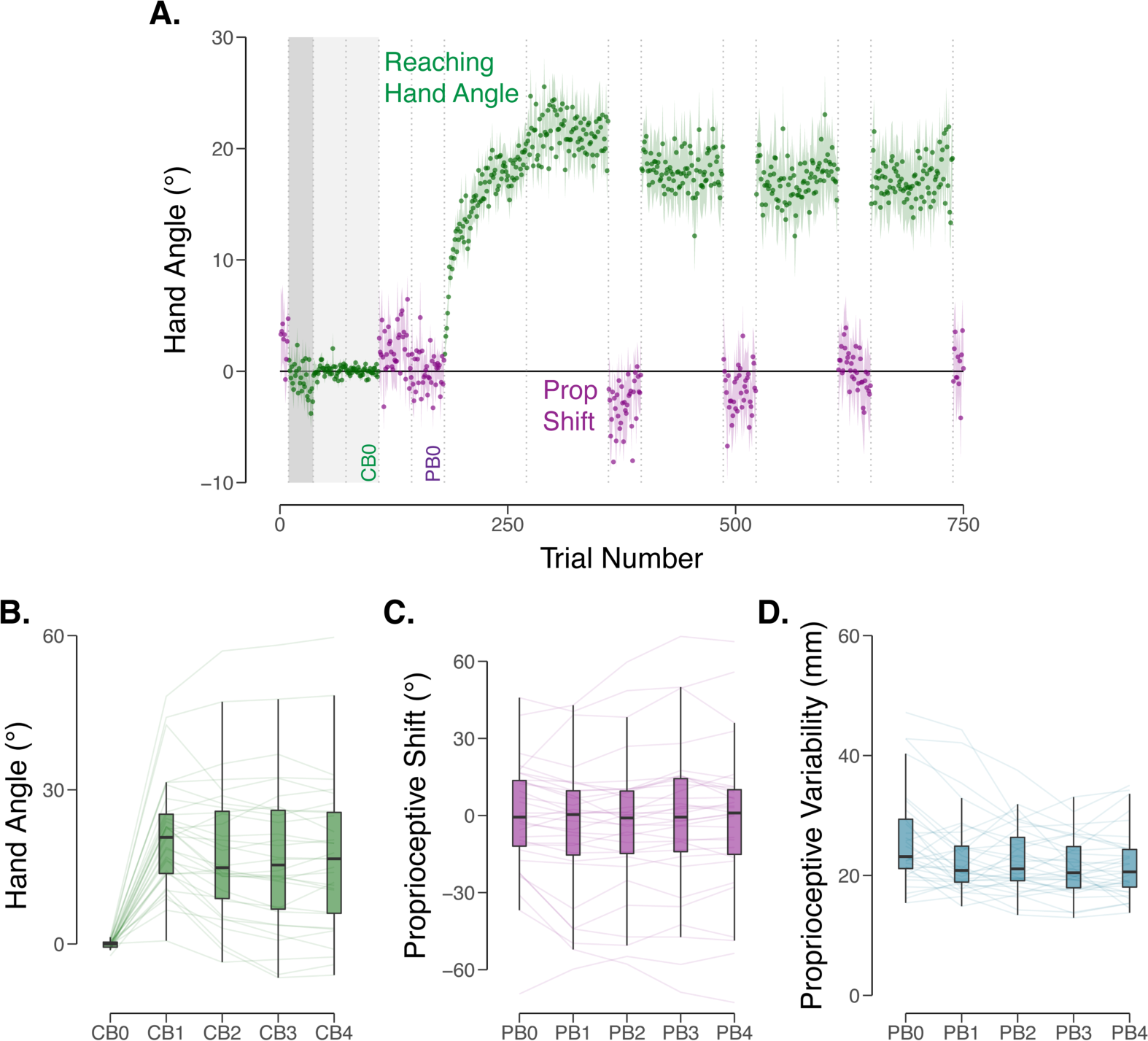
Performance on adaptation and proprioception probe tasks in experiment 2. **A)** Group means across test session. After a period of no feedback (dark grey region) and veridical feedback (light grey region) baseline trials, participants were exposed to a visual clamp in which the feedback was offset by 15° from the target. Participants performed blocks of reaching trials (hand angle shown in green) and proprioceptive probe trials (shift in perceived position shown in purple). Vertical dotted lines indicate block breaks. Shaded regions indicate ± SEM. Baseline blocks for reaching hand angle (CB0) and proprioceptive probes (PB0) are labeled. **B)** Mean hand angle averaged over the last three clamped feedback blocks. **C)** Proprioceptive error for each proprioceptive block. Thin lines indicate individual subjects. Box plots indicate min, max, and the 1^st^/3^rd^ interquartile range. **D)** Variability of proprioceptive judgments for each proprioceptive probe block.

#### Proprioceptive Measures

The proprioceptive shift in Exp 2 was modest, and in fact, only marginally significant (*F*_4,127_= 2.17, *p* = 0.08, *η*^2^ = 0.06). The mean value was -1.2° (sd = 11.37°), a mean that was smaller than the mean value of -4.0° (sd = 3.2°) observed in Exp 1 (*t*_59,_ = –4.3, *p* < 0.001, *d* = –1.1). 12 of the 32 participants exhibited a shift in the direction opposite to the cursor (compared to 2 out of 30 in Exp 1). Not only was the between-subject variability larger, but we also observed a large increase in Exp 2 in the within-subject variability of the proprioceptive reports (Exp 1, day 1: 21.7 ± 0.5, range = 12.3 – 36.0, Exp 2: 26.1 ± 8.0, range = 15.4 – 47.2; *t*_53_ = 2.5, *p* = 0.01, *d* = 0.6).

The large increase in the variability of the proprioceptive judgments is especially puzzling given that the two experimental protocols are very similar. It is possible that the clamped, non-contingent feedback used in Exp 2 has a different impact on sensed hand position compared to the contingent feedback provided in Exp 1. Alternatively, it may be related to other methodological differences. In particular, the studies were run by different experimenters, and they may have differed in how they passively displaced the participant’s arm, perhaps moving at different speeds.

Nonetheless, the proprioceptive shift and proprioceptive variability scores remained relatively stable across Exp 2. As noted above, in terms of mean values, there was no effect of block for proprioceptive shift. There was an effect of block on proprioceptive variability (*F*_4,124_ = 11.7, *p* < 0.001, *η*^2^ = 0.08), with post-hoc t-tests showing that the variability was larger on the baseline block compared to subsequent blocks (all *t*_124_ = 3.68, *p* < 0.04, *d*_*z*_ > 0.65), similar to that observed in Exp 1. More important in terms of the correlational analyses reported below, individual differences were maintained across the blocks for both proprioceptive shift (all pairwise correlations following the introduction of clamped feedback, from PB1 – PB4: *R* > 0.87, *p* < 0.001) and proprioceptive variability (all pairwise correlations between PB0 – PB4: *R* > 0.76, *p* < 0.001).

#### Correlating Adaptation and Proprioception

The correlational analysis between the three dependent variables yielded a similar pattern as that observed in Exp 1 (Figure 6). Consistent with the multisensory integration hypothesis, there was a positive correlation between the asymptote of implicit adaptation and proprioceptive variability (*R* = 0.37, *p* = 0.035). Consistent with the proprioceptive realignment hypothesis, there was a negative correlation between the asymptote of adaptation and the magnitude of proprioceptive shift (*R* = –0.62, *p* < 0.001). There was no correlation between proprioceptive shift and proprioceptive variability (*R* = –0.06, *p* = 0.76).

**Figure 6.**
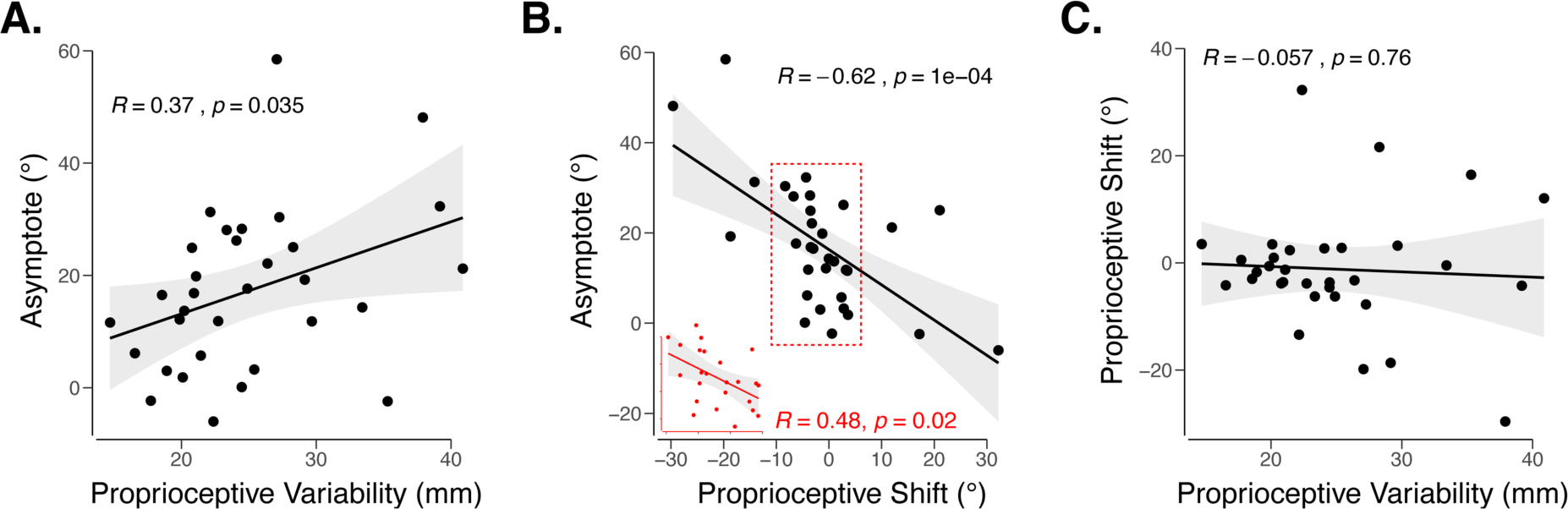
Inter-individual differences analyses in experiment 2. **A)** Proprioceptive variability vs asymptote in response to the visual clamp. **B)** Proprioceptive shift vs asymptote. A second correlation was performed on non-outlier data points contained in the red rectangle, also shown in the inset. **C)** Proprioceptive variability vs proprioceptive shift. Black line denotes the best fit regresion line and the shaded region indicates the 95% confidence interval.

We note that the correlations with the proprioceptive shift must be qualified. First, the effect of proprioceptive shift was only marginally significant. Second, there were extreme values in both directions, including participants who showed a large shift in the opposite direction of the expected shift (i.e., away from the clamped feedback). As a more conservative estimate, we repeated the correlational analyses using a non-parametric Spearman correlation (*R* = –0.57, *p* < 0.001), after removing all positive shifts (Figure 6B inset: 20 out of 32 individuals remaining, *R* = –0.70, *p* < 0.001), a subset of extreme positive shifters (4 individuals with more than 10° shift removed, 28 out of 32 individuals remaining: *R* = –0.72, *p* < 0.001) or a subset of extreme shifters in either direction (9 individuals with more than ±10° shift removed, 23 out of 32 individuals remaining: *R* = –0.48, *p* = 0.02). The correlation, in all cases, remained significant, evidence for a robust relationship between proprioceptive shift (albeit small) and implicit adaptation.

## DISCUSSION

The sensorimotor system uses visual and proprioceptive feedback to remain properly calibrated. Recent sensorimotor adaptation studies using visual perturbations to induce recalibration have revealed an upper bound on this process, beyond which changes in performance require alternative learning processes. While the contribution of vision to adaptation has been well characterized (Burge et al., 2008; Wei & Körding, 2010), the contribution of proprioception to adaptation remains poorly understood. Here, we took an individual differences approach, asking whether the extent of adaptation is correlated with biases and/or variability in the perceived position of the hand during adaptation.

There were two key findings: First, participants with greater proprioceptive variability in both experiments exhibited more implicit adaptation, a finding consistent with the multisensory integration account (Burge et al., 2008; Ernst & Banks, 2002; Robert J. van Beers, 2012; Wei & Körding, 2010). The asymptotic level of adaptation, in this view, reflects an equilibrium between learning from visual and proprioceptive error signals. This finding is consistent with adaptation being driven by the optimal weighting of proprioception and vision according to their relative variability, whereby greater proprioceptive variability results in greater weighting of visual feedback, and thus greater implicit adaptation. Second, participants with larger proprioceptive shifts towards the visual feedback exhibited larger implicit adaptation, a finding consistent with the proprioceptive realignment hypothesis (see also, Cressman & Henriques, 2009, 2010; Ruttle et al., 2020; Salomonczyk et al., 2013). The asymptotic level of adaptation, in this view, reflects the point of realignment between the sensed hand position and the target.

### Proprioceptive variability and asymptotic adaptation

Greater proprioceptive variability predicted a greater asymptotic magnitude of implicit adaptation. While we are unaware of any prior reports of this positive correlation, a recent study asked a related question: Does proprioceptive variability predict the early learning rate in response to the abrupt introduction of a 30° visuomotor rotation (Lei & Wang, 2018). This study reported no correlation between proprioceptive variability and early learning in young adults and a negative correlation in older adults. While these observations may appear inconsistent with the results of our study, their main dependent variable, early learning, likely reflects a strong contribution from explicit processes in response to this large perturbation (Bond & Taylor, 2015; Haith et al., 2015; Werner et al., 2015), rather than implicit adaptation. By this view, the null result for the young adults would suggest that proprioceptive variability is not related to explicit learning, whereas the negative correlation observed in older adults may reflect a concurrent age-dependent deterioration of strategy use and proprioceptive acuity (Vandevoorde & Orban de Xivry, 2019). Interestingly, older adults have also been shown to exhibit an age-dependent boost in implicit adaptation (Vandevoorde & Orban de Xivry, 2019). By the multisensory integration hypothesis, this increase would be expected if a decline in proprioceptive sensitivity is accompanied by an increase in proprioceptive variability, a hypothesis that can be tested using an individual difference approach in an older adult sample.

Previous tests of the multisensory integration account of implicit adaptation have focused exclusively on manipulations of the visual feedback. Increasing visual variability, either by replacing a small cursor with a cloud of dots or a Gaussian blur, has been shown to decrease the rate and extent of implicit adaptation (Burge et al., 2008; Körding & Wolpert, 2004; Tsay, Avraham, et al., 2020; Wei & Körding, 2010). Surprisingly, the sensory integration models put forth to account for these effects have not measured proprioception; rather, this component has either been estimated as a free parameter or ignored entirely.

Here we obtained direct measures of proprioceptive variability to test a core prediction of the multisensory integration model. A limitation with our individual difference approach, however, is that the analyses are purely correlational. Future studies using experimental methods to perturb proprioception (e.g., tendon vibration) (Bernier, Chua, Inglis, & Franks, 2007; Gilhodes, Roll, & Tardy-Gervet, 1986; Goodwin, McCloskey, & Matthews, 1972; Manzone & Tremblay, 2020; Roll, Gilhodes, & Tardy-Gervet, 1980) could build on our results, asking whether proprioceptive variability has a causal role in modulating the upper bound of adaptation.

### Proprioceptive shifts and asymptotic adaptation

In line with the proprioceptive realignment hypothesis, the upper bound of implicit adaptation was also correlated with the proprioceptive shift induced by the visual perturbation: Larger shifts were associated with (a negative correlation because of the direction used to measure the shift). We note that the proprioception realignment hypothesis offers an alternative multisensory integration perspective on adaptation, albeit one that entails *two* distinct processes that involve the integration of weighted signals. One integration process is the biased sense of hand position that arises with the introduction of the perturbed visual feedback, resulting in a proprioceptive shift. The size of the shift presumably reflects the relative weighting of the expected hand position (i.e., at the target) and the attractive force of the visual feedback. The second integration process involves the proprioceptive shift and the actual hand position. The weighted sum of these two signals defines the error signal that drives adaptation. Thus, as the hand adapts in the opposite direction of the target, the signal from the actual hand position can eventually negate the (stable) proprioceptive shift. The current results would suggest that the proprioceptive shift is given much more weight than the actual hand position: In the group means, a proprioceptive shift of ∼3° is only offset when the hand has adapted to around ∼20°.

Verbal reports of sensed hand position obtained in a continuous manner *during* adaptation provide converging evidence of the dynamics predicted by the proprioceptive realignment hypothesis. The report data followed a striking non-monotonic function, initially biased towards the clamped cursor (away from the target), and then reversing direction (Tsay et al., 2020). However, the asymptotic value of the reports was not at the target. Rather, it was shifted slightly away from the target in the opposite direction of the clamp. This “overshoot” is not predicted by either the multisensory integration or proprioceptive realignment hypotheses, a puzzle that remains to be addressed in future research.

The correlation between proprioceptive shift and implicit adaptation was only significant in Exp 2, although a similar the pattern was observed in Exp 1. One notable difference between the two experiments is the perturbation schedule: The rotation was introduced gradually in Exp 1, whereas the clamped rotation was introduced abruptly in Exp 2. This pattern is similar to that reported in a series of studies by Salomonczyk and colleagues. They used a force channel to move the hand along a trajectory that was deviated from the target by 30° while the participants saw a visual cursor move directly to the target. In this condition, a strong correlation was observed between the induced proprioceptive shift and adaptation (Salomonczyk et al., 2011, 2013). In contrast, when a 30° rotation of the cursor was gradually introduced (similar to Exp 1), the correlation was reduced by 50% and no longer significant (Salomonczyk et al 2011).

The difference between the abrupt and gradual conditions in terms of relating adaptation to changes in proprioceptive shift may be reconciled by the notion of causal inference, where the motor system is thought to infer the cause and therefore the relevance of different sources of sensory feedback (Berniker & Kording, 2011; Wei & Körding, 2009). When *contingent* visual feedback is provided (Exp 1 in the present study and Salomonczyk et al., 2011), the discrepancy between the position of the target and perturbed feedback not only signals an SPE, but also signals a task error—the cursor is missing the target. Given that task success requires adjusting the hand direction to make the cursor intersect the target, the sensed hand position may be deprioritized; thus, the sensed hand position would be given less weight and have a smaller impact on implicit adaptation. In contrast, when the visual feedback is *not contingent* on the movement and rendered un-informative (Exp 2 in the present paper), the sensed hand position may be a primary input for sensorimotor recalibration.

### Reconciling multisensory and proprioceptive realignment hypotheses

The core predictions for both the multisensory integration and proprioceptive realignment hypotheses were confirmed in the present experiments. These results motivate the following question: Do these two hypotheses, one based on the variability of proprioception and the other based on the shift in proprioception, reflect the operation of distinct processes involved in implicit sensorimotor adaptation? The absence of a correlation between the two proprioceptive measures, a finding consistent with several previous reports (Block & Bastian, 2011; Cressman & Henriques, 2009; Izawa, Criscimagna-Hemminger, & Shadmehr, 2012; Zaidel et al., 2011), is consistent with a dual-process model (see also, Block & Bastian, 2011). By this view, the observed asymptote is a composite of these two forms of adaptation. That is, a ∼20° asymptote is actually an equilibrium point between one process that weights the visual and proprioceptive inputs and a second process that seeks to nullify the proprioceptive shift.

Alternatively, there may be a more complex interaction between processes sensitive to proprioceptive variability and bias. We could envision a multi-stage process in which a reliability weighting rule operates at an early stage, whereas the integration of multiple inputs arises in a non-weighted manner at a later stage (or vice versa). Examples of the former are found in the optimal integration literature (Körding et al., 2007; Shams & Beierholm, 2010; Takahashi, Diedrichsen, & Watt, 2009; Wei & Körding, 2009). Examples of the latter are also ubiquitous, where proprioception and vision interact in a fixed manner, independent of variability (Rand & Heuer, 2019, 2020; Zaidel, Ma, & Angelaki, 2013; Zaidel et al., 2011). These stages of processing result in a final error signal, one that ultimately drives motor adaptation. While these ideas remain to be fleshed out in future research, the current results underscore the critical role of proprioception in sensorimotor adaptation.

## Author Contributions

All authors contributed to the study design. Testing, data collection, and data analysis were performed by D.E.P, A.R.S, and J.S.T. All authors contributed to the interpretation of results under the supervision of R.B.I. All authors drafted the manuscript and approved the final version of the manuscript for submission.

## Acknowledgements

This work was supported by grants R35 NS116883, R01 NS105839, and R01 NS1058389 from the National Institutes of Health (NIH). H.E.K. was funded by grants K12 HD055931 from the NIH and M3×1934650 from the National Science Foundation.

